# Role of transmembrane spanning domain 1 in cystic fibrosis transmembrane conductance regulator folding

**DOI:** 10.1101/2021.02.02.428178

**Authors:** Anna E. Patrick, Linda Millen, Philip J. Thomas

## Abstract

Cystic fibrosis (CF) is caused by mutations in the cystic fibrosis transmembrane conductance regulator (CFTR) protein that disrupt its folding pathway. The most common mutation causing CF is a deletion of phenylalanine at position 508 (ΔF508). CFTR contains five domains that each form cotranslational structures that interact with other domains as they are produced and folded. CFTR is comprised of two transmembrane spanning domains (TMDs), two nucleotide binding domains (NBDs) and a unique regulatory region (R). The first domain translated, TMD1, forms interdomain interactions with the other domains in CFTR. In TMD1, long intracellular loops extend into the cytoplasm and interact with both NBDs via coupling helices and with TMD2 via transmembrane spans (TMs). We examined mutations in TMD1 to determine the impact on individual domain and multidomain constructs. We found that mutations in a TM span or in the cytosolic ICLs interfere with specific steps in the hierarchical folding of CFTR. TM1 CF-causing mutants, G85E and G91R, directly affect TMD1, whereas most ICL1 and ICL2 mutant effects become apparent in the presence of TMD2. A single mutant in ICL2 worsened CFTR trafficking in the presence of NBD2, supporting its role in the ICL2-NBD2 interface. Mutation of hydrophobic residues in ICL coupling helices tended to increased levels of pre-TMD2 biogenic intermediates but caused ER accumulation in the presence of TMD2. This suggests a tradeoff between transient stability during translation and final structure. NBD2 increased the efficiency of mutant trafficking from the ER, consistent with stabilization of the full-length constructs. While the G85E and G91R mutants in TM1 have immediately detectable effects, most of the studied mutant effects and the ΔF508 mutant are apparent after production of TMD2, supporting this intermediate as a major point of recognition by protein quality control.

## Introduction

Cystic fibrosis (CF) is a common, lethal monogenetic disease caused by mutations in the cystic fibrosis transmembrane conductance regulator protein (CFTR) [1–3]. CFTR acts as a chloride channel at the surface of secretory epithelia, and its loss of function results in a reduced liquid layer and thick mucus secretions [4]. CF-causing mutations are identified throughout the protein, however 90% of CF patients have at least one allele with a deletion of phenylalanine at position 508 (ΔF508) [2]. The ΔF508 and other mutations result in misfolding of the CFTR protein, ER accumulation, and eventually degradation. Thus, the majority of CF is caused by mutations that disrupt CFTR folding.

CFTR is a member of the ABC transporter superfamily that uses ATP binding and hydrolysis to move substrates across membranes [5]. CFTR contains two transmembrane spanning domains (TMDs) and two nucleotide binding domains (NBDs) like other ABC transporters. CFTR also has a unique and less structured regulatory region (R) [6]. ABC transporter structures have varying resolutions and include Sav1866 [7, 8], MsbA [9], P-glycoprotein [10], MRP1 [11], zebrafish CFTR [12], and human CFTR [13]. Several homology models of CFTR have been constructed [14–17]. In CFTR, two TMDs wrap around each other in a domain-swapped fashion with each having two long intracellular loops (ICLs) that extend into the cytoplasm and interact with both NBDs. The NBDs interact in a head-to-tail fashion sandwiching the ATP binding sites [18–20]. Conformational signals from NBD ATP binding and hydrolysis are transmitted by the ICLs to the TMDs, resulting in chloride channel opening and closing [21, 22]. A coupling helix at the end of each ICL interacts with the NBDs. The coupling helices of ICL1 and ICL3 interact with both NBDs, with primary interactions occurring between ICL1 and NBD1 or ICL3 and NBD2. The coupling helices of ICL2 and ICL4 interact with NBD2 and NBD1 respectively [14].

As CFTR is translated, each domain folds and can then interact with previously translated domains to form multidomain intermediates [23–29]. The domains are translated in the order: TMD1-NBD1-R-TMD2-NBD2. A minimal construct of TMD1-NBD1-R-TMD2 is able to traffic from the ER [24, 27, 29, 30]. Finally, NBD2 undergoes a posttranslational incorporation into the CFTR structure [24]. A mutation in one domain can destabilize other domains in CFTR, highlighting the importance of interdomain interactions in this process [29, 31]. In this study, we use specific mutations in TMD1 to test this process.

The first domain, TMD1, contains extracellular loops, six transmembrane spans (TMs), and cytosolic regions including an N-terminal region, two ICLs, and a linker that connects to NBD1. The surface formed between the ICL1 coupling helix and NBDs is a mix of hydrophobic and hydrophilic interactions, whereas the surface formed by ICL2 is largely hydrophobic [14]. During TMD1 translation, regions that later form interdomain interactions are present for minutes until interaction partners are produced. Whether ICL structure forms in the TMD1 biogenic intermediate is not known.

CF-associated mutations are within the TMs and ICLs of TMD1 (www.genet.sickkids.on.ca). Many of these mutations introduce charge into the hydrophobic TM spans, and are thereby predicted to disrupt TMD1 structure [28]. In TM1, the CF-causing G85E and G91R mutations result in accumulation of CFTR in the ER [32]. These mutants maintain normal CFTR TM1 topology and have multidomain constructs with reduced stability [33]. The G85E mutant destabilizes the TM1 span in the ER membrane, producing an early defect for cellular recognition [32]. It is not clear whether G85E and G91R cause cellular recognition of TMD1 alone or if this requires multidomain intermediates.

CF-associated mutations in ICL1 and ICL2 are mostly in the helical bundle with few near the coupling helices (www.genet.sickkids.on.ca)[34]. For comparison, ICL4 interacts near the F508 position and contains far more CF-mutations [35, 36]. Predicted ICL1 and ICL2 amino acid residues include 141 to 194 and 242 to 307 respectively [14]. These sequences contain four disease-causing mutations that inhibit maturation of CFTR (H139R, G149R, D192G, and R258G) and two that reduce CFTR function (G178R and E193K) [34]. Specific residues are predicted as critical for ICL1 and ICL2 interactions with NBDs, including Y275 and W277 with NBD2 [17, 37] and D173,S169, and R170 with nucleotide and NBD1 [17]. These mutations have not been experimentally tested in multidomain folding intermediates.

The ΔF508 mutation alters multiple steps during CFTR folding. The ΔF508-NBD1 domain has tends to aggregate, is destabilized, and folds less efficiently, indicating a domain folding problem [26, 38–41]. NBD1 structure forms during translation [25] and ΔF508 effects are apparent during translation [24, 41]. In full-length ΔF508-CFTR crosslinks between the TMDs and NBDs are altered [15, 42] and multidomain folding intermediates are destabilized [24, 27, 29, 31, 43]. The ΔF508 mutant effects can be rescued independently by suppressor mutations within NBD1 [41, 44–46] and by a suppressor mutation in ICL4 [47]. Both these steps have additive effects on rescue of the ΔF508 mutant protein [48]. The additive effects on rescuing and potentiating ΔF508 mutant protein is essential for therapeutics used to treat CF patients [49]. Importantly, current clinical therapeutics stabilize TMD1 [50, 51]. Understanding the role of TMD1 in CFTR folding is important for ongoing therapeutic development.

This study examines biogenic intermediates and full-length CFTR containing CF-causing mutations in TM1, basic residue mutations in the ICL1 and ICL2 coupling helices, and the ΔF508 mutation to test their roles in CFTR domain and multidomain folding.

## Materials and methods

### Plasmids, DNA techniques

An expression plasmid of full-length, wild type CFTR (pCMV-CFTR-pBQ4.7) was a gift from J. Rommens (The Hospital for Sick Children, Toronto) and was mutagenized using standard protocols for site-directed mutagenesis [52]. Site-directed mutagenesis was performed by PCR techniques using PfuUltra high-fidelity DNA Polymerase (Stratagene). All mutations were confirmed by DNA sequencing. In these constructs, two stop codons were introduced after amino acid residues 414, 670, or 1174. For TMD1 C-terminal truncations, stop codons were introduced after positions 414, 388, and 354 in wild type or G85E mutant CFTR generated from the pBQ4.7 CFTR cloned into a pBI bidirectional vector. An ECL1 glycosylation site in CFTR is previously described [32]. Stop codons were introduced into this construct to make truncation mutants.

### Multiple sequence alignment

ABC transporters of the exporter type include Sav1866 (pdb 2ONJ and 2HYD), MsbA from E. coli (pdb 3B5W), MsbA from V. cholerae (3B5X), MsbA from S. typhimurium (3B60), P-glycoprotein from mouse (pdb 3G5U, 3G60, 3G61). A multisequence alignment was by entering the UniProt accession numbers for these proteins into ClustalW and selected regions examined. UniProt accession numbers are human CFTR (P13569), Sav1866 (Q99T13), MsbA from *E. coli* (P60752), *V. cholerae* (Q9KQW9), and *S. typhimurium* (P63359), and P-glycoprotein from mouse (P21447). The coupling helices for CFTR were selected based on the identified helices in the highest resolution structure, Sav1866 [7].

### Mammalian cell protein expression

HEK293 cells (American Type Culture Collection, ATCC) were maintained in Dulbecco’s Modified Eagle Medium (DMEM, Invitrogen) supplemented with 10% Fetal Calf Serum (Gemini Bio-Products), 50μg mL^−1^ penicillin, and 50 units mL^−1^ streptomycin using standard culture techniques. CFTR constructs were transfected at 20,000 cells per mL in suspension using polyetheylenimine, (PEI Polysciences Inc.) transfection reagent, plated in 24 well plates, and expressed for 48 hours. Cell lysis was performed in RIPA buffer (20mM tris pH7.6, 150mM NaCl, 0.1% SDS, 1% IGEPAL, 0.5% deoxycholic acid, 1mg EDTA-free protease inhibitor tablet (Roche)) at 4°C and centrifuged at 13000g’s to generate cleared lysate. Sample buffer (60mM tris pH6.8, 5% glycerol, 2% SDS, bromophenol blue, 280mM β-mercaptoethanol) was added to cleared lysate and incubated at 37°C for 20 minutes.

### Western blot analysis

SDS-PAGE analysis of full-length CFTR was performed on equal volumes of lysate. Samples were transferred to PVDF Immobilon membranes (Millipore) and Western blotting performed using CFTR antibodies or actin antibody (Millipore). The CFTR antibody utilized for full-length protein was 596 against amino acids 1204-1211 (UNC School of Medicine), for TMD2x was 570 against amino acids 731-742 (UNC School of Medicine), and for TMD1x, NBD1x, and Rx was MM13-4 (AbCAM). The secondary antibody for all primary antibodies was peroxidase-AffiniPure goat anti-mouse IgG (Jackson ImmunoResearch Laboratories, Inc.). All Western blots were developed using Amersham ELCPlus Western blotting detection reagent (GE Healthcare).

### Quantitation

Signal from all membranes was collected by film and using a Typhoon 9410 Variable Mode Imager (GE Healthsciences). The TMD1x, NBD1x, and Rx samples were quantified from film and the TMD2x and full-length samples were quantified from Typhoon. Quantification was performed using Image J Software. Each membrane contained all samples, and individual sample intensity was normalized to total membrane sample intensity. This was then compared across membranes.

### Glycosylation analysis

Glycosylation analysis of the ECL1 site was performed on HEK293 cells transiently transfected with the ECL1 site constructs, expressed for 48 hours, and lysed in RIPA buffer. 500U of PNGaseF (New England Biolabs) or Endoglycosidase H (New England Biolabs) was added to 40μL of cleared lysate and incubated at 37° for 2 hours. Sample buffer was added to the reaction and incubated at 37° for 20 minutes before SDS-PAGE and Western blot analysis using antibody MM13-4 (AbCAM). For each transfected construct, mock treated lysate was loaded immediately next to EndoH and PNGaseF treated lysate to detect gel shifts that occur upon glycosylation removal.

## Results

### Mutations in TMD1 alter CFTR cellular trafficking

TMD1 is an integral membrane domain and has amino acid residues in the cytosol, ER membrane, and ER lumen during CFTR translation. Positions in TM1, ICL1, and ICL2 known to disrupt CFTR folding or with predicted locations near the ICL coupling helices were perturbed by introduction of charge. The positions selected are mapped onto the homologous Sav1866 structure (Fig 1A). The CF-causing mutations, G85E and G91R, are in TM1. The G178K, M265R, and W277R CF-associated mutations are in ICL1 and ICL2. These mutations are predicted to introduce a basic residue close to the coupling helices (www.genet.sickkids.on.ca). The coupling helices for CFTR were modeled by aligning the sequences of homologous exporter structures and selecting residues in CFTR utilizing the Sav1866 structure (Fig 1B). The mutated positions within CFTR and respective positions in other transporters are highlighted (Fig 1B). Positions were mutated to the basic amino acid residues lysine or arginine.

**Fig 1.**
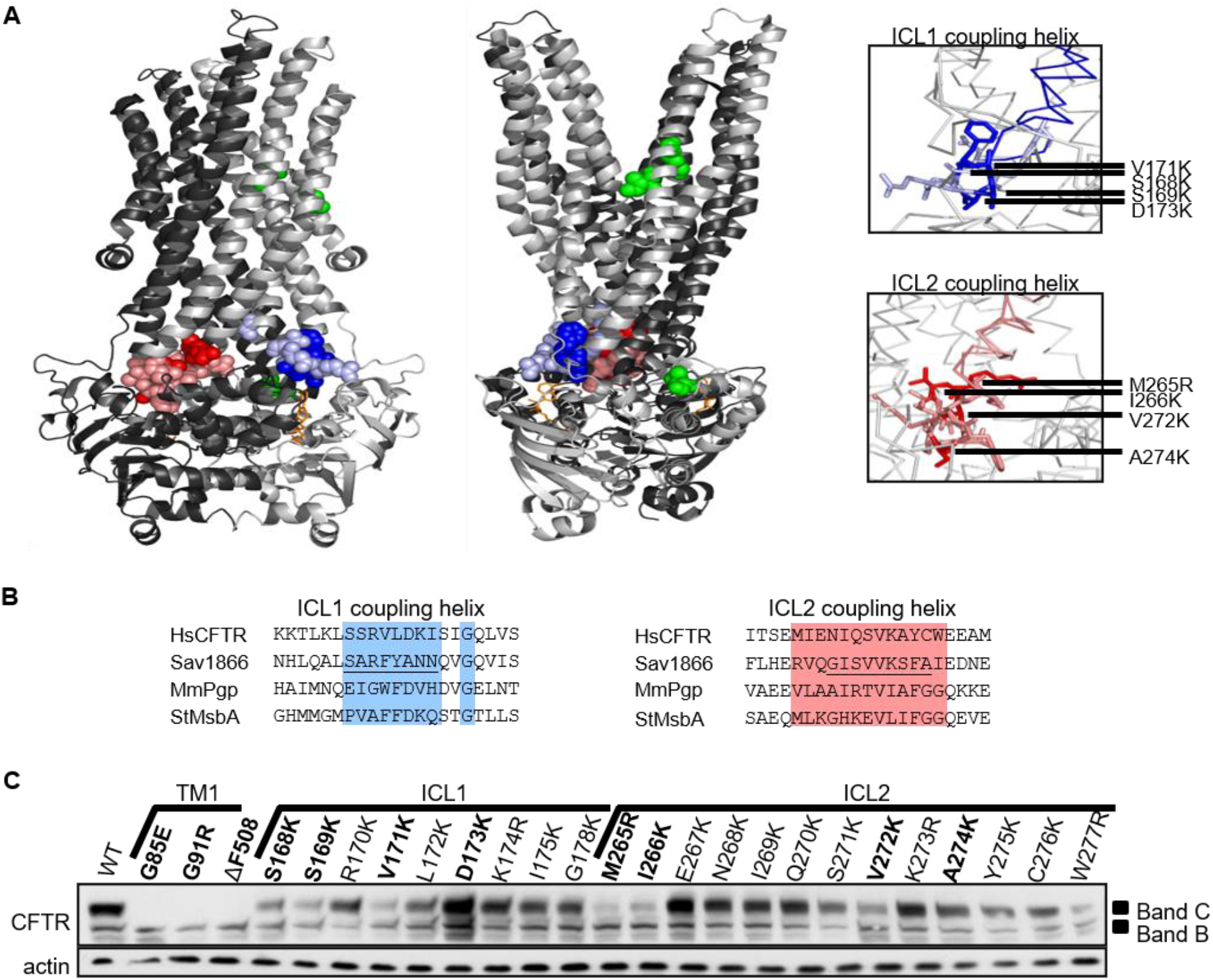
TMD1 mutant effects on full-length CFTR cellular trafficking. (A) Mutations mapped onto the Sav1866 structure (pdb 2HYD). Sav1866 is a homodimer, and each protein unit is colored with light gray representing CFTR TMD1 and NBD1 and dark grey representing CFTR TMD2 and NBD2. Respective CFTR mutation positions are shown as spheres for TM1 and ΔF508 (green), ICL1 (light blue), and ICL2 (light red). Mutants that were further studied are a darker blue or red. Views down ICL1 and ICL2 coupling helices axes with the protein backbone as a ribbon and mutated residues as sticks. (B) Alignments of human CFTR with ABC exporters using ClustalW software and sequences for *H. sapiens* CFTR, *S. aureus* Sav1866, *M. musculus* P-glycoprotein, and *S. typhimurium* MsbA (UniProt accession numbers: P13569, Q99T13, P21447, and P63359). Screened CFTR mutant positions and respective positions in transporters are highlighted blue for ICL1 and red for ICL2. The coupling helix for Sav1866 is underlined. (C) TMD1 mutants examined in full-length CFTR. Expression and trafficking in HEK293 cell culture with assessment by Western blot analysis using antibody 596. Band B is core glycosylated in the ER and Band C has been complex glycosylation in the Golgi. Mutants further studied are bolded. The ΔF508 mutation accumulates in the ER and is a control for Band B. Actin is a loading control.

The TM1, ICL1, and ICL2 mutations were introduced into full-length CFTR, transiently transfected in HEK293 cell culture, and CFTR maturation monitored by Western blot analysis (Fig 1C). The progression of CFTR as it traffics in the cell is reflected by the glycosylation status of two natural N-linked glycosylation sites in TMD2 that become core glycosylated in the ER (Band B) and are then modified by complex glycosylation in the Golgi (Band C). The ΔF508 mutation accumulates in the ER as Band B. The CF-causing TM1 mutants G85E and G91R accumulate in the ER, indicated by the sole presence of Band B. Mutations at the test positions are integrated into the ER and mature at a variety of efficiencies as compared to WT. G85E, G91R, ΔF508, S168K, S169K, V171K, D173K, M265R, I266K, V272K, and A274K were chosen for further study based on inefficient maturation or predicted location in the ICL-NBD interfaces.

### Generation of CFTR biogenic intermediates

We made CFTR multidomain biogenic intermediates to examine TMD1 interactions with other domains. To generate biogenic intermediates, stop codons, designated by x, were introduced after different domains based on the structural model, resulting in constructs producing residues 1-414 (TMD1x), 1-670 (NBD1x), 1-836 (Rx), and 1-1174 (TMD2x) (Fig 2A). Consistent with other studies [27, 29], the full-length and TMD2x constructs traffic from the ER and are complex glycosylated in the Golgi (Fig 2B). The TMD1x, NBD1x, and Rx constructs do not contain the natural N-linked glycosylation sites. Membrane integration and trafficking of these constructs was monitored using a reporter glycosylation site introduced in extracellular loop 1 [32]. The TMD1x, NBD1x, and Rx constructs are core glycosylated, but not complex glycosylated, indicating integration into the ER membrane but inefficient trafficking to the Golgi (Figs 2B and 2C).

**Fig 2.**
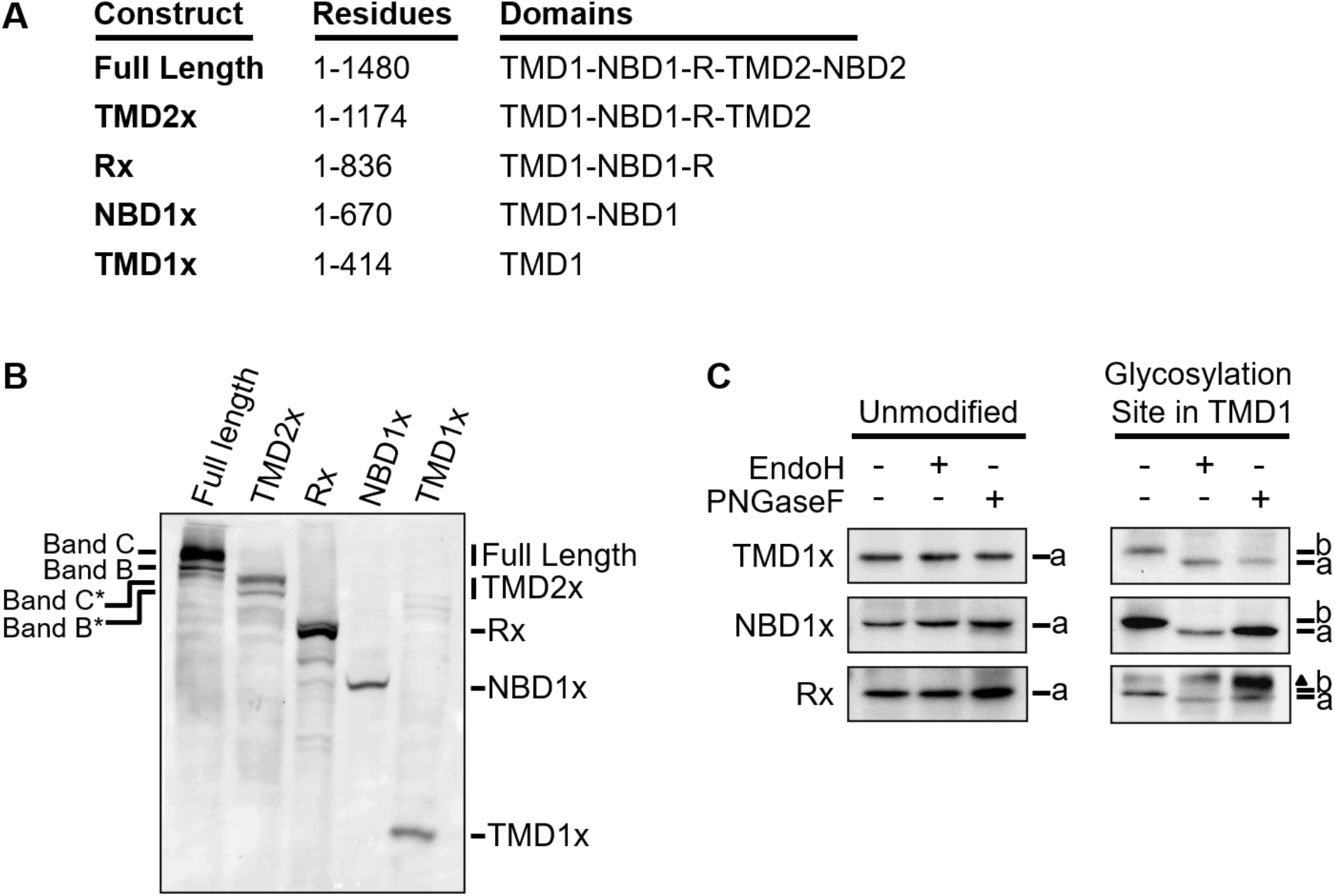
CFTR biogenic intermediates. (A) The CFTR biogenic intermediates, residues, and multidomain construct produced by introducing stop codons. (B) Full-length CFTR (1-1480) and biogenic intermediates were transiently expressed in HEK293 cells and assessed by Western blot analysis with mixed CFTR antibodies 570 and MM13-4. TMD2 contains the two natural glycosylation sites for monitoring cellular trafficking of CFTR. Full-length Bands B and C and 1174x core glycosylated Band B* and complex glycosylated Band C* are indicated. (C) A reporter core glycosylation site introduced into TMD1 assessed in TMD1x, NBD1x, and Rx. Unmodified biogenic intermediates or biogenic intermediates with the TMD1 glycosylation site were expressed in HEK293 cells and examined by treatment with glycosidase followed by Western blot analysis with MM13-4 antibody. Glycosidase treatment used EndoH to remove core glycosylation and PNGaseF to remove both core and complex glycosylation. The positions of core glycosylated (b) and non-glycosylated (a) constructs are noted. Unrelated background in the TMD1 glycosylation site Rx sample is designated by a triangle.

TMD1 forms cotranslational structure that then can interact with other domains or proteins. The C-terminal boundary of TMD1 was examined by making constructs containing different C-terminal residues. Stop codons were introduced after residues 354 (354x), 388 (388x), and 414 (414x) to include all TM6, an additional linker region, and the N-terminal residues of NBD1 respectively (Fig 3A). Steady state levels of transiently transfected constructs in HEK293 cells were examined by Western blot analysis for wild type and G85E mutant protein (Fig 3B). The 354x construct expresses at low levels for both wild type and G85E, suggesting this construct is less stable (Fig 3B). The wild type 388x and 414x constructs have higher steady-state levels compared to the G85E 388x and G85E 414x constructs (Fig 3B). This suggests that G85E effects are identifiable at steady state levels in TMD1 constructs containing more than 388 residues.

**Fig 3.**
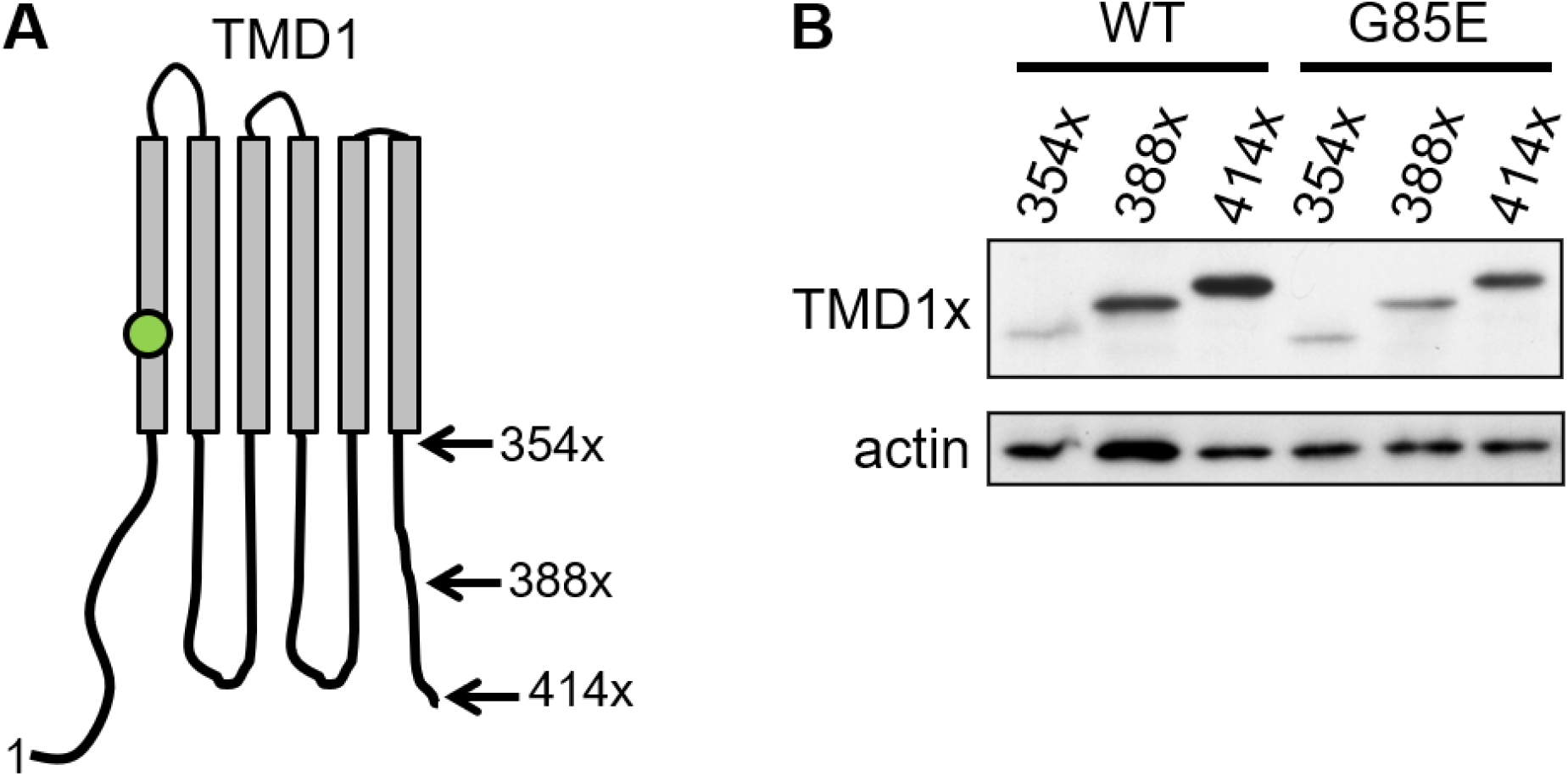
TMD1 C-terminal boundary mapping. (A) A schematic of the TMD1 C-terminal stop codon placement after TM6 (354x), after the linker region (388x), or including N-terminal NBD1 residues (414x) is depicted with the position of G85E indicated (green circle). (B) Wild type (WT) and G85E constructs were transiently transfected into HEK293 cells and monitored by Western blot analysis with the antibody MM13-4. Actin is a loading control.

### TMD1 mutant impact on TMD1x

Selected TMD1 mutations were examined in the TMD1x biogenic intermediate containing residues 1-414 (Fig 4A). Steady state levels of transiently transfected constructs in HEK293 cells were examined by Western blot analysis, quantified, and normalized to total image CFTR signal (Fig 4B). The CF-causing mutants G85E and G91R have significantly reduced steady-state levels with respect to wild type. The mutant D173K replaces an acidic with a basic residue and has a significantly decreased level with respect to wild type. Mutations that replace a polar residue with a basic residue, S168K and S169K, had no effect on steady-state TMD1x. Many mutants that replace a hydrophobic residue with a positively charged residue (V171K, I266K, V272K, and A274K) have an increased steady-state level of TMD1x. Importantly, the mutant effects in TMD1x are different than in full-length, indicating other domains are required for their full impact.

**Fig 4.**
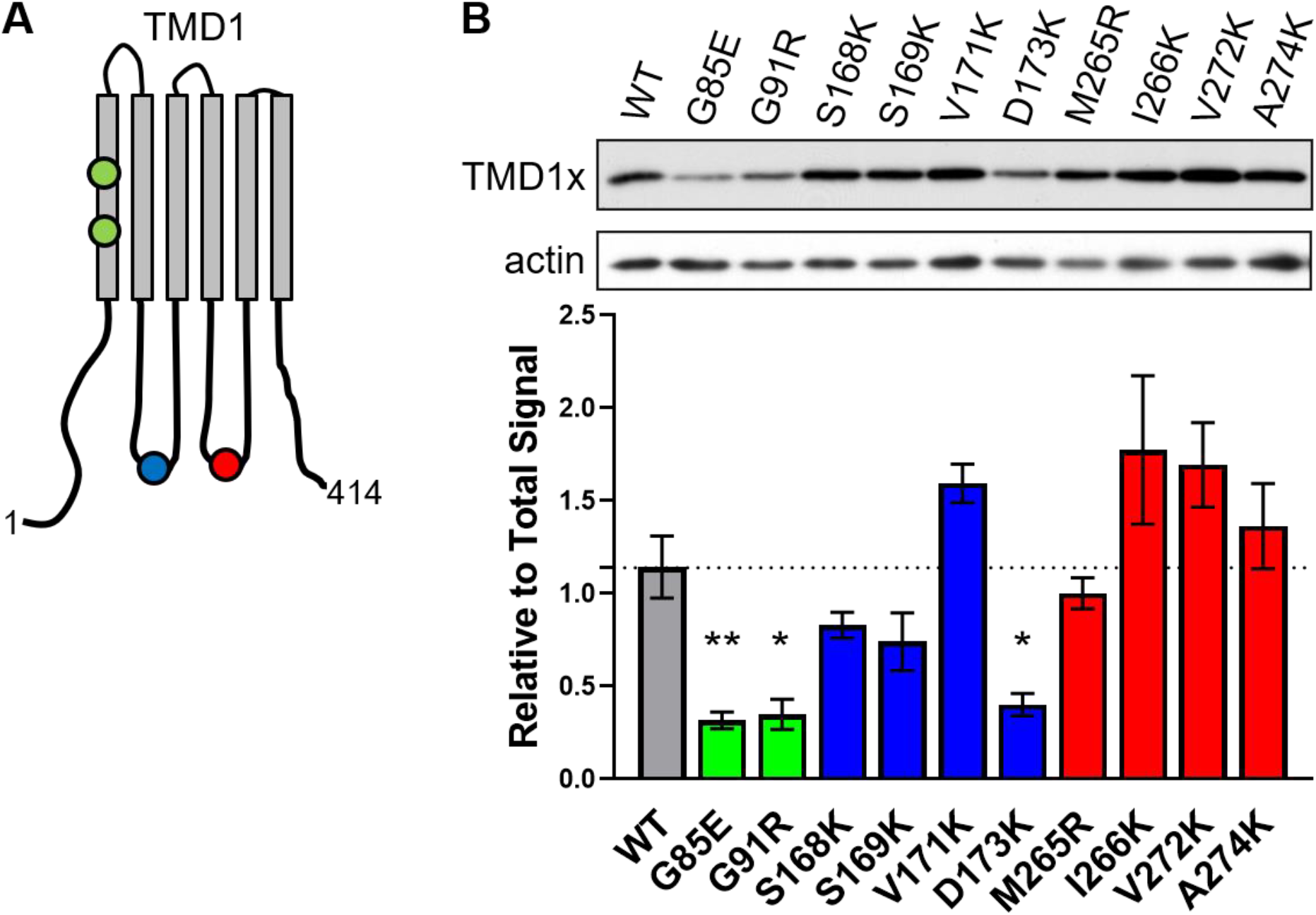
TMD1 mutants in the TMD1x biogenic intermediate. (A) Schematic of TMD1x (residues 1-414) with six transmembrane spans, ER luminal, and cytosolic regions. Approximate locations of mutants in TM1 (green circles), ICL1 (blue circle), and ICL2 (red circle) are marked. (B) TMD1x containing a mutation was transiently transfected in HEK293 cells and monitored by Western blot analysis with the antibody MM13-4. The steady-state levels were quantified and normalized to total image TMD1x signal with the average and standard error of the mean plotted (N=4). TM1 mutants are green, ICL1 mutants are blue, and ICL2 mutants are red. Actin is a loading control. Analysis by ANOVA against WT, * p<0.05, ** p<0.01.

### TMD1 mutant effects are less apparent in NBD1x and Rx

The TMD1 mutants were studied in the NBD1x construct. The NBD1x construct contains residues 1-670 of CFTR (Fig 5A). In this construct, mutations can disrupt an individual domain or interdomain interactions. Steady-state levels of the constructs were monitored after transient transfection in HEK293 cells by Western blot analysis (Fig 5B). The CF-mutant G85E significantly reduces steady state levels as compared to wild type, indicating NBD1 does not rescue nor amplify the defect. G91R and D173K trend to a lower expression, however unlike the TMD1x construct this is not statistically significant. Again, multiple mutants that replace a hydrophobic residue with a positively charged residue tend to increase steady-state levels of NBD1x. Most mutants have NBD1x levels the same as wild type, suggesting that the cell maintains steady state levels of NBD1x for wild type and most mutants in a similar fashion.

**Fig 5.**
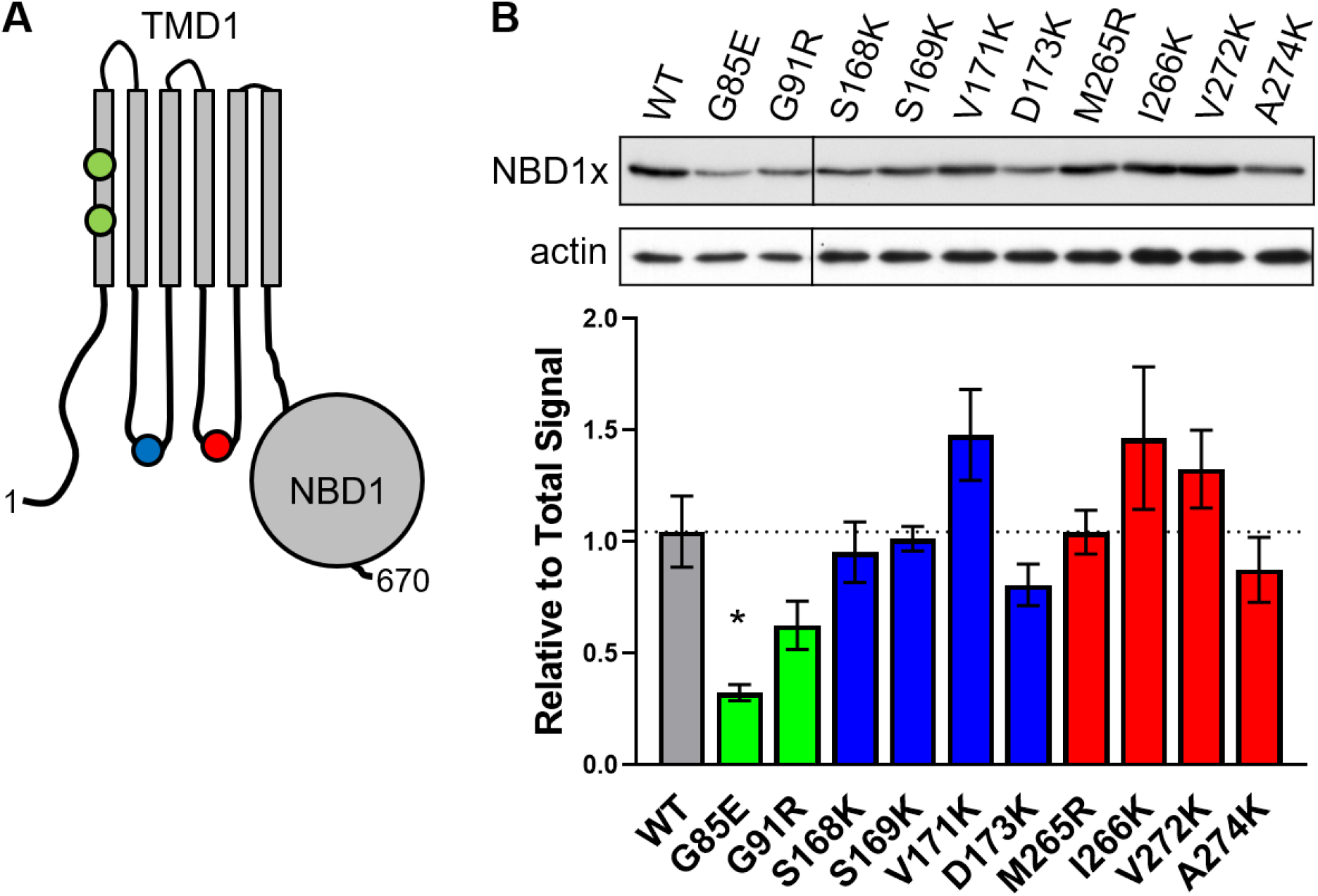
TMD1 mutants in the NBD1x biogenic intermediate. (A) Schematic of NBD1x construct (residues 1-670) with TMD1 and NBD1. Approximate locations of mutants in TM1 (green circles), ICL1 (blue circles), and ICL2 (red circles) are marked. (B) NBD1x containing a mutation was transiently transfected into HEK293 cells and monitored by Western blot analysis with the antibody MM13-4. The steady-state levels were quantified and normalized to total image NBD1x signal with the average and the standard error of the mean plotted (N=4). TM1 mutants are green, ICL1 mutants are blue, and ICL2 mutants are red. Actin is a loading control. Analysis by ANOVA against WT, * p<0.05.

The R domain has potential interaction surfaces for a biogenic intermediate. Other studies found that CFTR constructs truncated after the R domain had a longer half-life than shorter constructs both in cell culture and *in vitro*, suggesting an increased stability [33, 43]. We examined the selected TMD1 mutants in the Rx construct containing residues 1-836 (Fig 6A). Steady state levels of no mutants were statistically different from wild type, thought a decreased trend is seen for G85E, G91R, and D173K (Fig 6B). Again, the mutants that replace a hydrophobic residue with a positively charged residue show a tendency to increase steady-state levels of Rx. Mutants have Rx levels more like wild type steady-state cellular levels than either the TMD1x or NBD1x biogenic intermediates.

**Fig 6.**
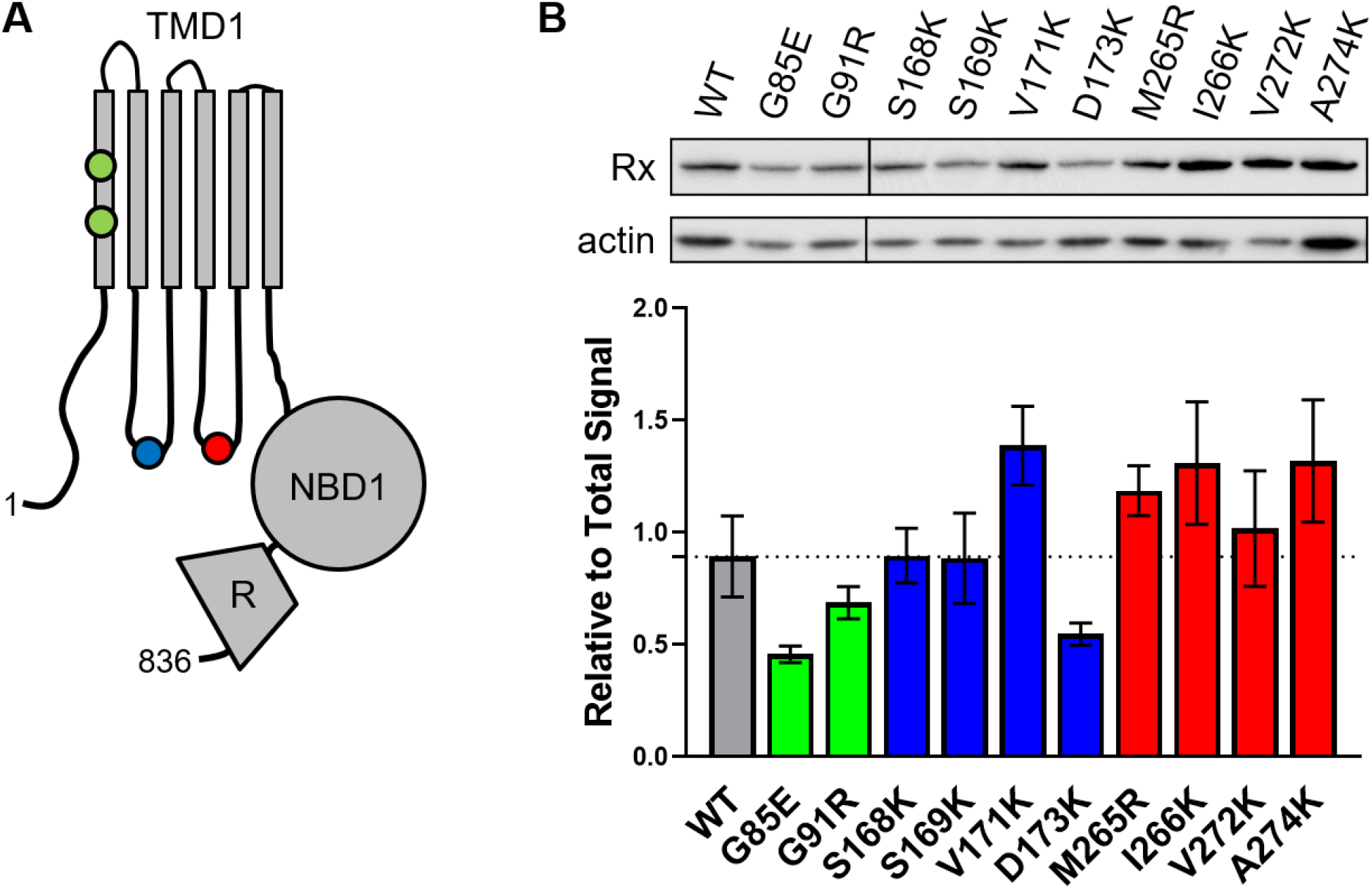
TMD1 mutants in the Rx biogenic intermediate. (A) Schematic of Rx construct (residues 1-836) with TMD1, NBD1, and R. Approximate locations of mutants in TM1 (green circles), ICL1 (blue circles), and ICL2 (red circles) are marked. (B) Rx containing a mutation was transiently transfected into HEK293 cells and monitored by Western blot analysis with the antibody MM13-4. The steady-state levels were quantified and normalized to total image Rx signal with the average and the standard error of the mean plotted (N=4). TM1 mutants are green, ICL1 mutants are blue, and ICL2 mutants are red. Actin is a loading control. Analysis by ANOVA against WT, * p<0.05.

### TMD1 mutant effects are dramatic in TMD2x

The TMD2x construct contains the first four domains, TMD1-NBD1-R-TMD2, and forms a trafficking competent structure that can be monitored by glycosylation of sites in TMD2 (Fig 2B). The TM1, ICL1, and ICL2 mutations were examined in TMD2x containing residues 1 to 1174 of CFTR (Fig 7A). The CF-causing mutants G85E and G91R accumulate in the ER in TMD2x indicated by presence of solely Band B* (Fig 7B). Most ICL mutants have a reduction in steady-state levels of trafficked TMD2x, with significant reduction in band C* (Figs 7B). This is a striking difference from the effect of TMD1x, NBD1x, and Rx constructs. This suggests that a stable TMD2x structure is disrupted by the TMD1 mutants, and this abnormal structure is recognized by the cell resulting in reduced steady-state levels. An exception is that D173K is like wild type, suggesting that the presence of TMD2 improved the significant defect in the TMD1x and NBD1x constructs. The A274K mutant remains like wild type. Interestingly, the V272K mutant shows a small but significant deviation from wild type that is less pronounced than the nearby M265R and I266K mutants. None of the mutants show a difference in the ER steady state levels of mutants as indicated by Band B* (Fig 7B).

**Fig 7.**
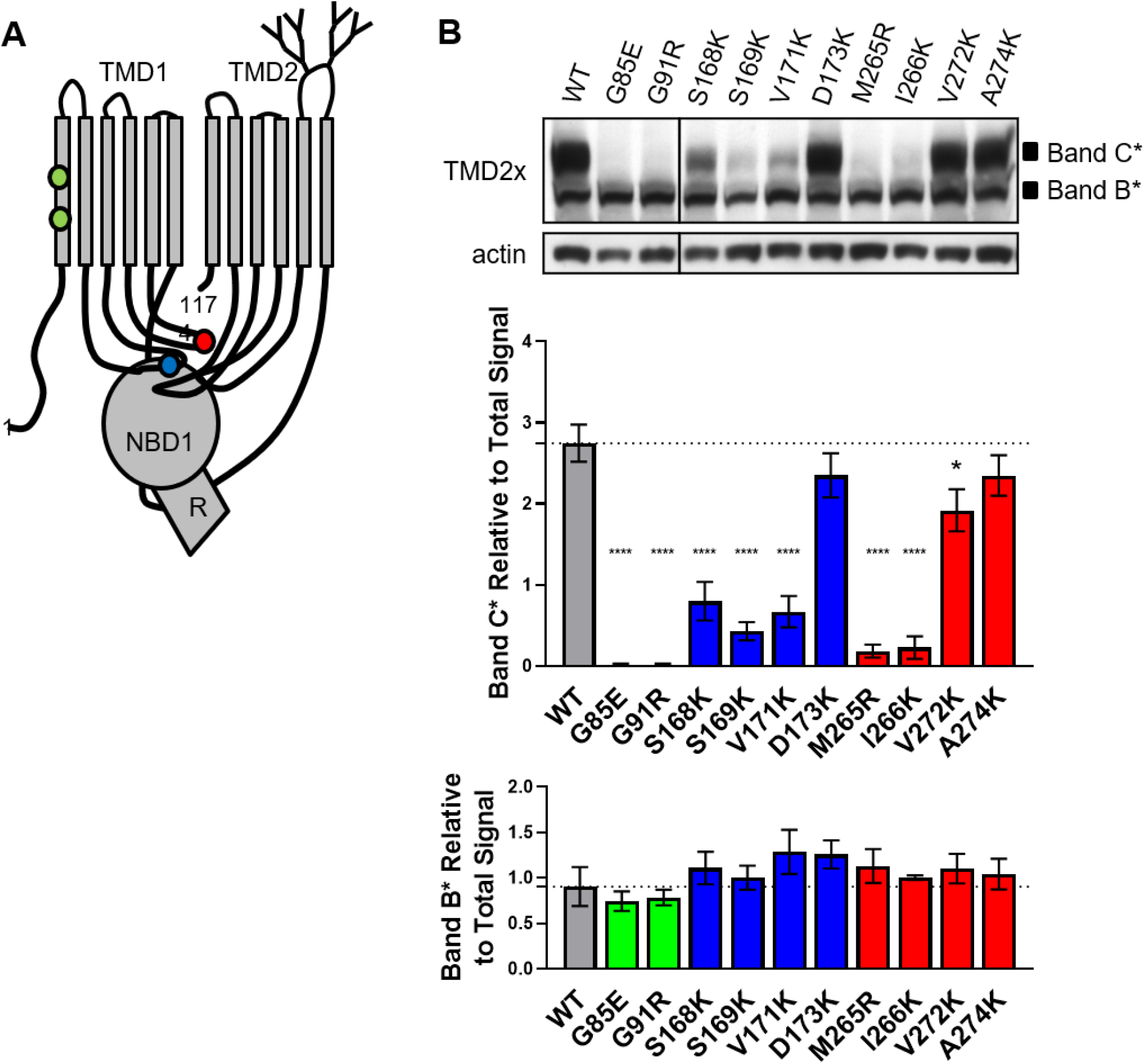
TMD1 mutants in the TMD2x biogenic intermediate. (A) Schematic of TMD2x (residues 1-1174x) with TMD1, NBD1, R, and TMD2. Approximate locations of mutants in TM1 (green circles), ICL1 (blue circles), and ICL2 (red circles) are marked. (B) TMD2x containing a mutation was transiently transfected into HEK293 cells and monitored by Western blot analysis with the antibody 570. CFTR steady-state levels of Band B* and Band C* were quantified and normalized to total image TMD2x signal. The average is plotted with the standard error of the mean (N=4). TM1 mutants are green, ICL1 mutants are blue, and ICL2 mutants are red. Actin is a loading control. Analysis by ANOVA against WT, * p<0.05, ** p<0.01, *** p<0.001, **** p<0.0001.

### NBD2 contributes to stabilization of CFTR

Full-length, functional CFTR contains extensive interactions between all five domains. Its cellular trafficking is monitored by glycosylation of sites in TMD2. The TMD1 mutants were examined in a full-length construct containing residues 1-1480 of CFTR (Fig 8A). The CF-causing mutants G85E and G91R accumulate in the ER without trafficking, indicated by a lack of Band C (Figs 8B). The D173K and A274K mutants have similar levels of Band C compared to wild type, which is like their effect in the TMD2x construct. In full-length, all other ICL mutants have a significantly reduced steady state level of trafficked construct, indicated by Band C. Importantly, the efficiency of ICL mutant trafficking is higher in the full-length protein than in TMD2x. This is consistent with NBD2 conferring stabilization and improved trafficking of CFTR. A notable exception is the V272K mutant, which exhibits a worse efficiency of trafficking in the full-length protein compared to TMD2x. The V272 position is centrally located within the coupling helix of ICL2, which forms major interactions with NBD2 that might only be disrupted in the full-length construct. In full-length constructs, a decrease in mutant trafficking from the ER is detected by a significant increase in Band B for mutants G85E, G91R, S168K, M265R, and I266K. Both mutants with wild type like trafficking, D173K and A274K, also have wild type like levels of Band B.

**Fig 8.**
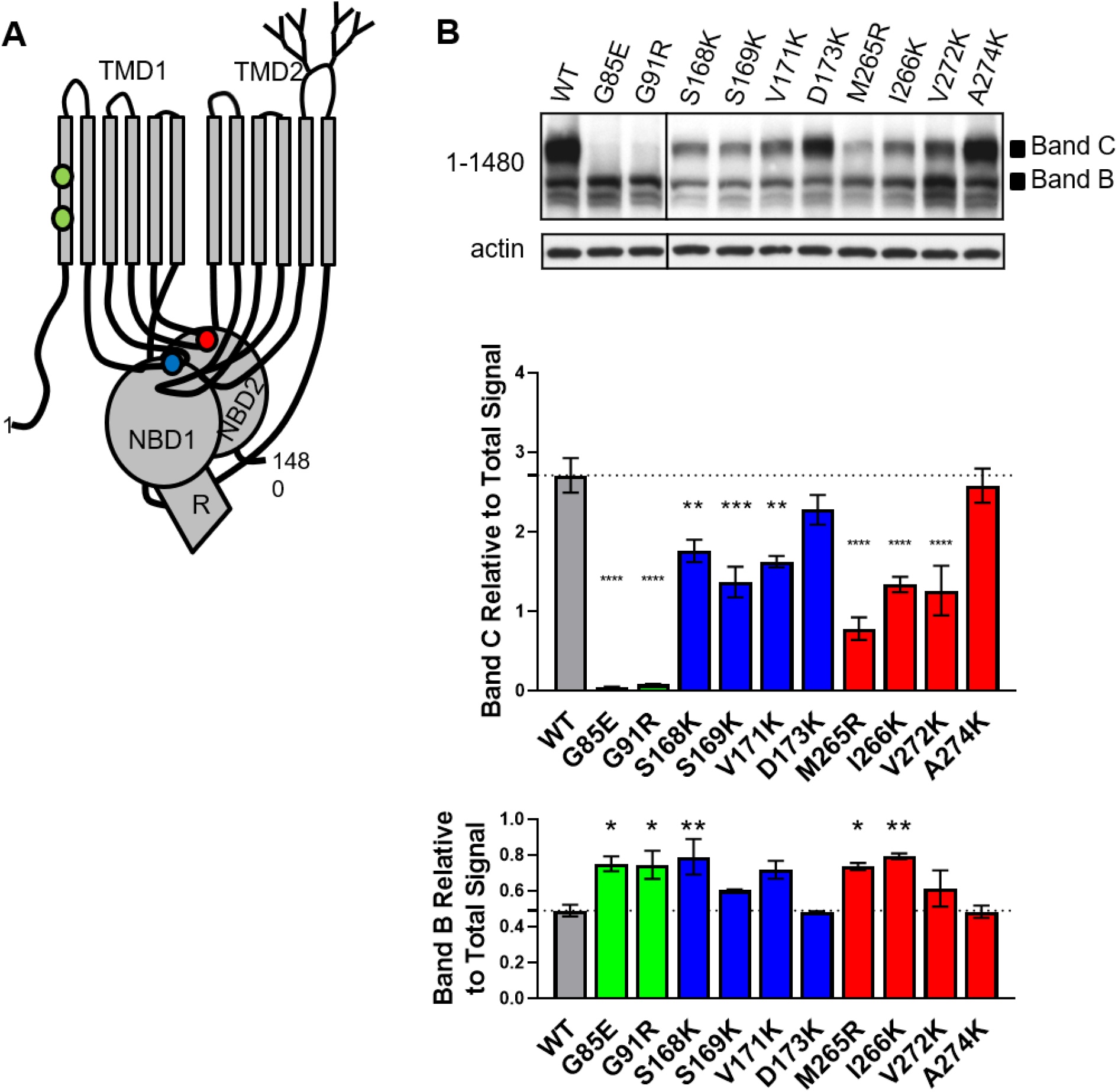
TMD1 mutants in full-length CFTR. (A) Schematic of full-length CFTR (residues 1-1480) depicts TMD1, NBD1, R, TMD2, and NBD2. Approximate locations of mutants in TM1 (green circles), ICL1 (blue circles), and ICL2 (red circles) are marked. (B) A full-length construct containing a mutant was transiently transfected into HEK293 cells and monitored by Western blot analysis with the antibody 596. CFTR steady-state levels of Band B and Band C were quantified and normalized to total image full-length signal. The average is plotted with the standard error of the mean (N=3). TM1 mutants are green, ICL1 mutants are blue, and ICL2 mutants are red. Actin was monitored as a loading control. Analysis by ANOVA against WT, * p<0.05, ** p<0.01, *** p<0.001, **** p<0.0001

### ΔF508 mutant effects are largely revealed in TMD2x

The biogenic intermediates containing the ΔF508 mutation were examined to determine the intermediate most significantly deviating from CFTR steady state levels. Multiple NBD1x constructs (n=9) and Rx constructs (n=7) were generated to minimize a plasmid specific effect. Constructs were transiently expressed in HEK293 cell culture in 6 biologic replicates, with multiple constructs in some experiments (Fig 9A). The steady-state levels of NBD1x ΔF508 is significantly reduced from wild type. The steady-state level of Rx ΔF508 is like wild type (Fig 9A). The ΔF508 mutant was examined in the TMD2x and full-length constructs. Constructs were transiently expressed in HEK293 cell culture and Bands B* and C* or Bands B and C quantified (Fig 9B). ΔF508 has dramatically lower steady-state levels of Band C* and Band C in TMD2x and full-length constructs respectively (Fig 9B). While early effects of ΔF508 are identifiable in NBD2x, the cell makes a major distinction between wild type and ΔF508 in TMD2x (Fig 9B). Similar to our other studies mutants, in TMD2x Band B* for ΔF508 is no different from wild type. In the full-length construct, ΔF508 has a significant accumulation of Band B in the ER (Fig 9B). A model depicts potential interactions between the domains of ΔF508-CFTR during translation with red arrows representing possible interdomain disruptions in the TMD2x and full-length constructs (Fig 9C).

**Fig 9.**
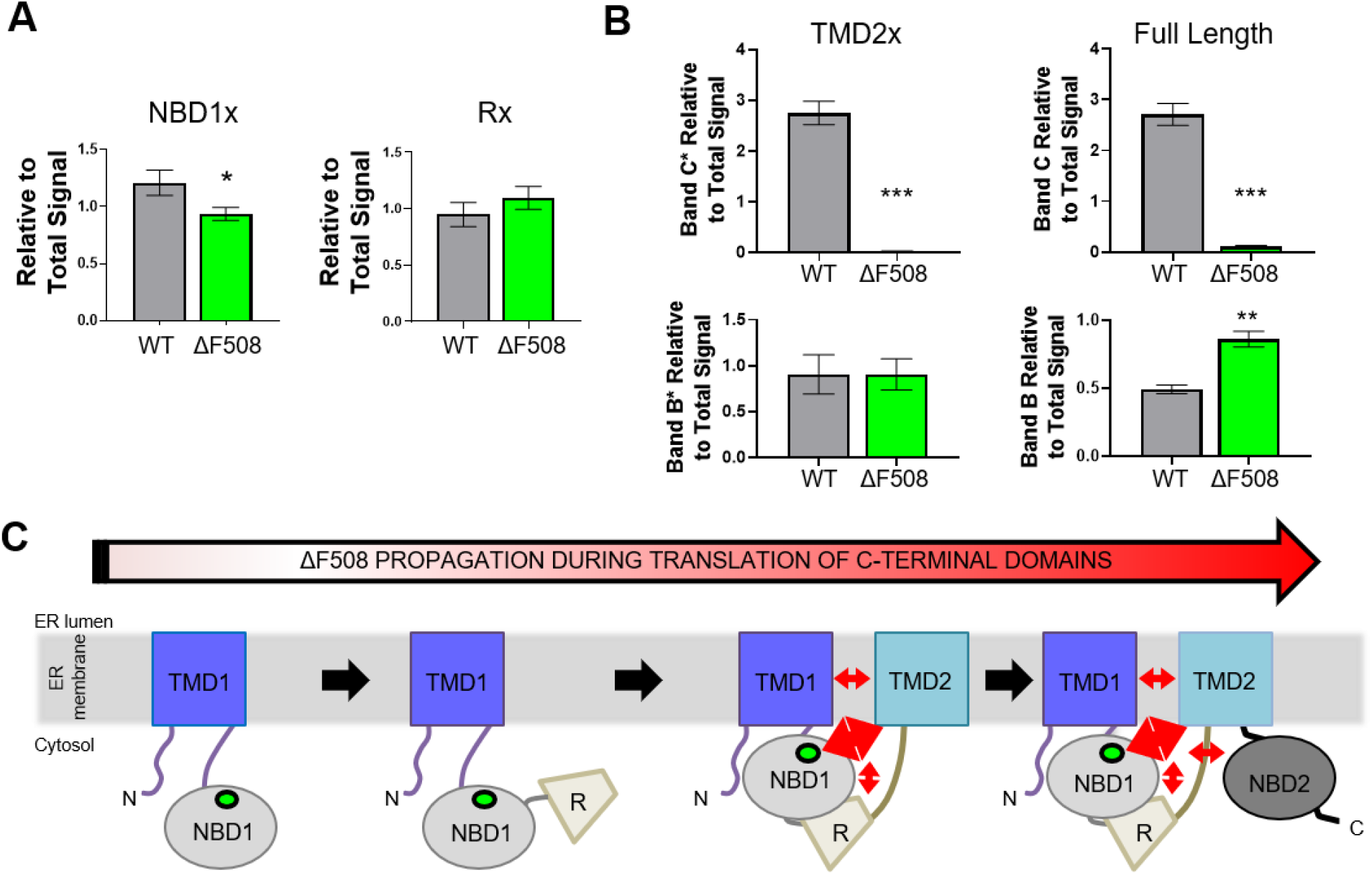
ΔF508 dramatically alters TMD2x and full-length CFTR. (A) Wild type (WT) and ΔF508 NBD1x and Rx constructs were transiently transfected into HEK293 cells and monitored by Western blot analysis. The average and standard error of the mean for NBD1x, WT (N=8) and ΔF508 (N=21), and Rx, WT (N=8) and ΔF508 (N=17), are from 6 biologic replicates. (B) WT and ΔF508 TMD2x and full-length constructs were transiently transfected into HEK293 cells and monitored by Western blot analysis. Steady-state levels of Bands B*, C*, B, and C were quantified and normalized to total image TMD2x and full-length respectively. The average is plotted with the standard error of the mean (N=3). (C) Depiction of a folding model with ΔF508 in the presence of different multidomain intermediates. Red arrows represent possible interdomain disruptions in the TMD2x and full-length constructs.

## Discussion

This study examines the role of TMD1 in formation of CFTR domain structure and interdomain intermediates. The structural units are disrupted by specific mutations in TMD1, including CF-causing mutations. CF-associated mutants and polymorphisms (www.genet.sickkids.on.ca) have been identified throughout TMD1. Furthermore, TMDs are stabilized by CF therapeutics. The corrector VX-809 (lumacaftor) stabilizes TMD1 [50] and also stabilizes TMD1 mutant CFTR and ΔF508-CFTR [51]. Understanding TMD1 and its role within the formation of CFTR structure is increasingly important since this domain is promising for future therapeutics.

Many disease-associated mutations introduce charge into TMD1 TM spans [28] or ICLs [34]. More CF-causing mutants have been identified in ICL1 than in ICL2. Of these, few are near the coupling helices suggesting that mutations in these regions do not result in CF or are not represented in the database. We found that introduction of basic residues in many positions in ILC1 and ICL2 did not completely block CFTR trafficking (Fig 1C). Thus, few individual residues in ICL1 and ICL2 have a major impact on the formation of the ICL-NBD interface. A similar finding occurs for growth hormone binding to the human growth hormone receptor, whereby few residue mutations in the protein interface alter binding affinity [53]. This results from the interface ability to alter binding energetics [54] and remodel [55] to maintain similar binding, which could relate to the described observations for CFTR ICL-NBD interactions. We identified selected mutations and studied them in biogenic intermediates. A model depicts the earliest biogenic intermediate wherein a persistent TMD1 mutant impact was found (Fig 10).

**Fig 10.**
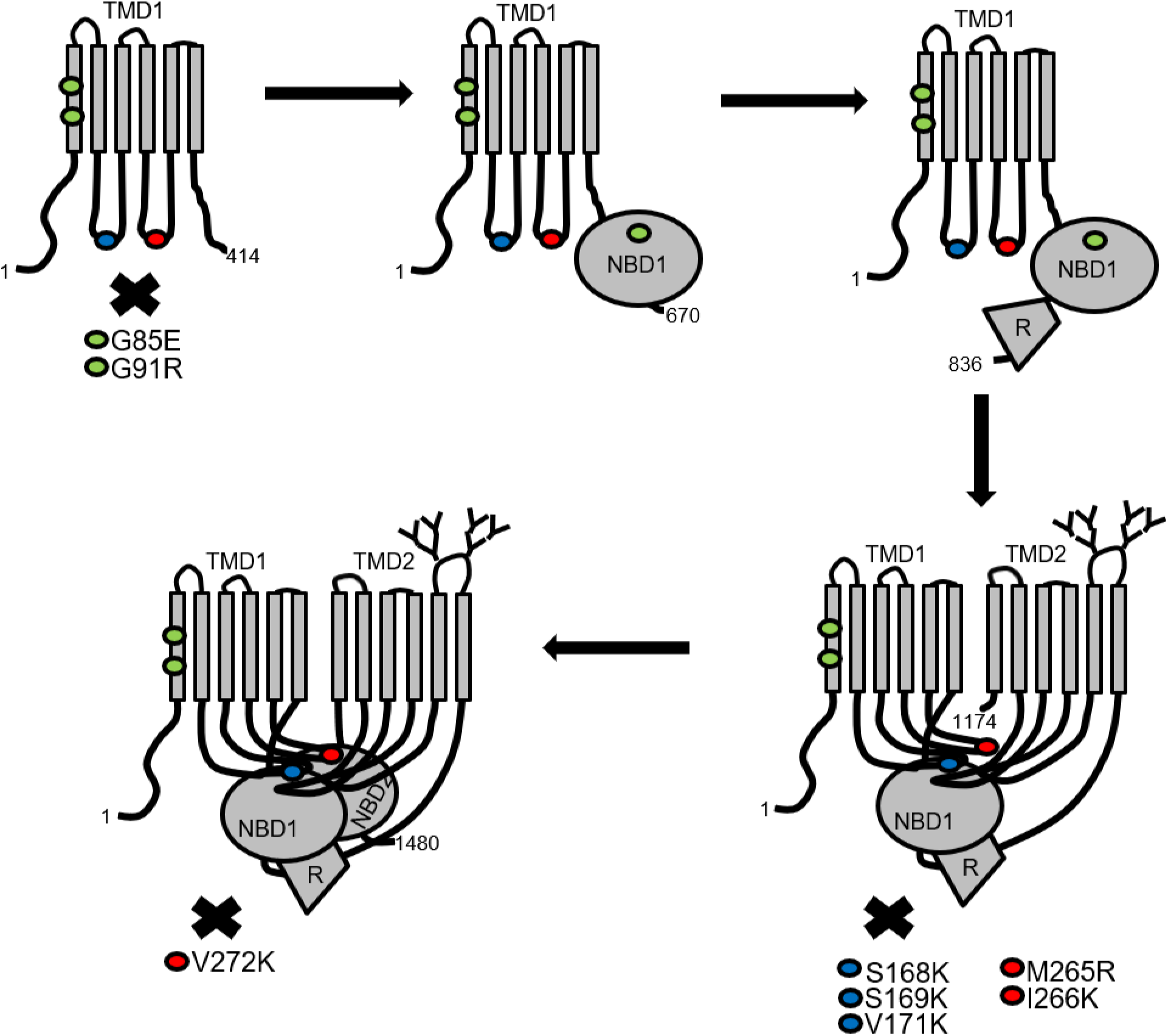
Model of TMD1 impact on biogenic intermediates. Schematic depicting the CFTR biogenic intermediates as they would occur during translation. Below each biogenic intermediate, an X denotes the initial intermediate in which the mutant has a significant and large impact on steady-state levels.

CFTR is monitored by protein folding and quality control proteins during translation [56]. Quality control involves a protein complex containing the E3 RMA1, E2 Ubc6e, and Derlin-1 interacting with N-terminal regions, and then the Hcs70/CHIP E3 complex interacting during and after more C-terminal regions are produced [43, 56–58]. TMD1 mutants could be recognized by either or both complexes. G85E mutant CFTR is recognized by the Derlin-1 associated cellular quality control machinery early during biogenesis [56, 59, 60]. Derlin-1 mediates the retro translocation and ER-associated degradation of misfolded proteins [61, 62]. Our study shows that G85E has effects within TMD1 (Fig 4). Our prior work showed that G85E destabilizes TM1 [63]. TMD1 likely contains structural signals for the recognition and degradation of the G85E mutant by quality control machinery. The G91R mutant is associated with the RMA1 E3 ubiquitin ligase that interacts with Derlin-1 [56]. G91R has effects within TMD1 (Fig 4). Yet, *in vitro* work suggests G85E and G91R are recognized in multidomain constructs [33]. For both mutants, it is likely that multiple levels of recognition are involved in cellular mistrafficking and reduced steady state levels.

TMD1 makes a proteolytically protected structure during translation [25]. Integration of the TM spans is required for its formation. If a TM span is marginally hydrophobic, the efficiency of integration becomes more efficient with flanking regions and other TM spans present [64]. In TMD1, TM6 has several basic residues that are important for pore formation [65]. TM6 *in vitro* integration is inefficient, and truncation at 364 results in TM6 incorrect integration [66]. In our study, we found that moving the end of TMD1 more C-terminal from 354x to 388x to 414x increased the steady state level of the TMD1x construct (Fig 3). In the presence of G85E, these construct levels were only reduced for 388x and 414x. This suggests that the 354x construct is not stable with reduced steady state levels and that G85E is not able to make this detectably lower. Multiple CF-associated mutants and polymorphisms are found in the region between 388-414 (www.genet.sickkids.on.ca). Our study supports an essential role for these C-terminal residues of TMD1 for stable domain formation and thereby CFTR function, which could be perturbed by mutations in this region.

ICL1 and ICL2 form extensive interdomain interactions in the final CFTR structure, yet their role during folding is not clear. In this study, we focused on the ICL coupling helices, which are cytosolic and approximately 25Å away from the membrane in homologous structures [7]. In CFTR the predicted coupling helix and NBD interfaces are both hydrophilic and hydrophobic for ICL1, and largely hydrophobic for ICL2 [14]. Surprisingly, the TMD1x, NBD1x, and Rx constructs trend towards increased steady-state levels when a hydrophobic residue is mutated to a basic residue. Thereby, the coupling helix hydrophobic residues may decrease level of biogenic intermediates. Hsc70 recognizes short hydrophobic sequences [67] and could form an interaction with the coupling helices. Furthermore, the Hsc70/CHIP machinery is found in association with TMD1 [43] and targets immature CFTR for degradation [43, 57, 58]. Our work supports a hypothesis that the coupling helices are monitored during CFTR folding through interactions with Hsc70, and persistent hydrophobic residue exposure can lead to targeting of misfolded CFTR for degradation.

The impact of most positively charged ICL mutants in TMD1 is detected in TMD2x (Fig 10). TMD2x contains domains wherein TMD rearrangements, ICL helical bundle structure, and interactions with NBD1 can occur. The TMD1 mutants have reduced trafficking efficiency from the ER, suggesting an intermediate that is recognized in the presence of TMD2 or failure of appropriate trafficking signals. A di-acidic motif within NBD1 is important for COPII-dependent trafficking of CFTR from the ER [68]. The exposure of this motif could depend on a structural rearrangement that occurs after TMD2 is produced. If TMD1 mutants do not have the native multidomain rearrangements, then this site may not be revealed resulting in mistrafficking.

In the full-length protein, most lysine mutants trafficked more efficiently than in the TMD2x construct. The addition of NBD2 confers a greater folding efficiency and trafficking from the ER. Thus, NBD2 is not strictly required for CFTR trafficking [27, 29, 30, 47] however its posttranslational association into the CFTR structure [24] imparts stability to the other domains. The exceptions to this are for the mutations D173K, A274K, and V272K. The D173K mutation in ICL1 reduces biogenic intermediates prior to TMD2 and behaves more like wild type in the presence of TMD2. One explanation is that this position is involved in formation of a nonnative contact that is lost in the constructs containing TMD2. The A274K mutation is like wild type in all constructs, suggesting this position is tolerant to mutation. The V272K mutant lowered trafficking efficiency in the full-length construct more dramatically than in TMD2x, which lacks NBD2. In CFTR homology model, position V272 points between the two helical stems of ICL2 to support ICL structure, thereby stabilizing the ICL2-NBD2 interface indirectly. An equivalent position in ICL4, L1065, has a CF-causing mutation of L1065P that accumulates in the ER [35, 36]. We propose that stabilization of the ICL and coupling helix structures rather than direct stabilization of the NBD/ICL interfaces should be considered a potential therapeutic target for improving mutant CFTR folding and function.

The major recognition of many CFTR mutants requires TMD2, which is also true for ΔF508 (Fig 9). Some reports indicate that ΔF508 minimally reduces stability of TMD1-NBD1 constructs, but significantly reduces stability of constructs containing the R domain [31, 69]. We found that the steady-state levels of NBD1x-ΔF508 is reduced from wild type and the Rx-ΔF508 construct is not altered from wild type. These data are inconsistent with previous reports. In our experimental data, we could observe this difference within the range of expressed individual constructs. We have tried to minimize this possibility by using multiple independently generated constructs in our studies between wild type and ΔF508. In TMD2x-ΔF508, the protein may fail to achieve a complex, multidomain fold that could allow the cell to distinguish misfolded from appropriately folded CFTR. The lower levels of NBD1x-ΔF508 may reflect misfolded domain, and the dramatic decrease in TMD2x-ΔF508 reflects disruption of interdomain interactions and multidomain structure formation. This is consistent with compounds or suppressor mutations for ΔF508 within NBD1 [44, 45, 70] or distant in the CFTR protein [71–73] improving trafficking and function. The importance of targeting multiple steps for ΔF508-CFTR is highlighted by clinical therapeutics used to treat CF patients [49].

This study identifies roles for TMD1 in CFTR folding and details for CF-causing mutant misfolding. This data is important for understanding CFTR misfolding and for identifying potential therapeutic targets in that process.

## Acknowledgments

We thank Juan Mendoza for help with sequence alignment and selection of the intracellular loop residues. We thank J. Rommens for providing the CFTR construct. This research was supported by NIH grants R37 DK049835 (P.J.T.) and K12 HD087023 (Research Scholar, A.E.P.).

